# Medium-throughput zebrafish optogenetic platform identifies deficits in subsequent neural activity following brief early exposure to cannabidiol and Δ-9-tetrahydrocannabinol

**DOI:** 10.1101/2021.01.04.425279

**Authors:** Richard Kanyo, Md Ruhul Amin, Laszlo F. Locskai, Danika D. Bouvier, Alexandria M. Olthuis, W. Ted Allison, Declan W. Ali

## Abstract

In the light of legislative changes and the widespread use of cannabis as a recreational and medicinal drug, delayed effects of cannabis upon brief exposure during embryonic development are of high interest as early pregnancies often go undetected. Here, zebrafish embryos were exposed to cannabidiol (CBD) and Δ-9-tetrahydrocannabinol l (THC) until the end of gastrulation (1-10 hours post-fertilization) and analyzed later in development (4-5 days post-fertilization). In order to measure neural activity, we implemented CaMPARI (Calcium-Modulated Photoactivatable Ratiometric Integrator) and optimized the protocol for a 96-well format complemented by locomotor analysis. Our results revealed that neural activity was decreased by CBD more than THC. At higher doses, both cannabinoids could dramatically reduce neural activity and locomotor activity. Interestingly, the decrease was more pronounced when CBD and THC were combined. At the receptor level, CBD-mediated reduction of locomotor activity was partially prevented using cannabinoid type 1 and 2 receptor inhibitors. Overall, we report that CBD toxicity occurs via two cannabinoid receptors and is synergistically enhanced by THC exposure to negatively impact neural activity late in larval development. Future studies are warranted to reveal other cannabinoids and receptors involved in this pathway to understand the subsequent health implications of cannabis consumption on fetal development.

## Introduction

Cannabis is consumed most commonly as a recreational drug that is often portrayed as harmless, yet the health implications are not fully understood^1^. The positive association of this drug in the public eye is not a surprise considering extracts of the *Cannabis sativa* plant have been used for medical purposes for almost 5000 years, especially for pain treatment^2^. However, scientific knowledge is limited and despite controversy, some countries, more recently Canada, continue to legalize cannabis for recreational use. Cannabis has been reported to be one of the most illicitly used drugs during pregnancy, with increase in consumption over the years, and key compounds, Δ-9-tetrahydrocannabinol (THC) and cannabidiol (CBD), can readily cross the placenta^3–8^. We were especially interested in CBD due to the reported health benefits and availability in many natural products including oils and food^9^. In the past, the negative impacts of cannabis in connection to embryonic development have been principally associated with the psychoactive THC^10^. In humans, epidemiological and clinical studies associate maternal cannabis exposure to behavioural disturbances in the offspring linked to increased risk for neuropsychiatric disorders^11^. In rats, maternal exposure of THC changed a series of behaviours in the offspring, including water-induced grooming, increased light sensitivity and altered exploratory behaviour^12^. Recently, the negative impacts of cannabis have expanded and also include the non-psychotropic CBD, which disrupts motor-neuron development in zebrafish^13^. This study is in contrast to reports that suggest positive health benefits of CBD, by treating nausea during pregnancy^14,15^.

The mechanistic pathway(s) by which THC and CBD are toxic are elusive. A new study linked these two cannabinoids to the sonic hedgehog signaling pathway in mice and zebrafish, and appears to involve cannabinoid type 1 receptor (CB1R)^16^. It is well established that THC binds and activates as a partial agonist two distinct classes of G-coupled protein receptors, CB1R and CB2R^17^. Both receptors are highly expressed in neuronal tissue where CB1R is localized mostly in the CNS^18–20^ and CB2R in the peripheral nervous system^21^, although CB2R has been also linked to the CNS^22–24^. CBD has been reported to interact with CB1R and CB2R, but with a lower affinity relative to THC, and rather antagonizes cannabinoid-induced effects indirectly through other receptors^25–29^. Additional data to illuminate signaling pathways of THC and CBD *in vivo* would highlight functional important mechanisms.

Zebrafish embryos have several advantages that complement mammalian models. Experiments are economical because many embryos are available, and exposure studies are simple as compounds can be easily added and removed. Further, embryos develop outside the female and therefore not confounded by maternal physiology and variable transport to fetus. Translucent larvae provide opportunities to implement cutting-edge fluorescing calcium sensors and measure neural activity. Past studies suggest that the endocannabinoid system plays a role in zebrafish development^30–32^. In addition, both cannabinoid receptors investigated here have similar expression profiles in the CNS compared to mammals, with CB1R sharing a 70 % protein sequence identity with the human homolog^33^. Therefore, the zebrafish is an excellent model organism to further illuminate biological mechanisms. Considering the anticipated effects of THC and CBD, high-throughput quantification of neural activity following early exposures would be of great interest. In this study, the calcium modulated photoactivatable ratiometric integrator (CaMPARI)^34^, was implemented. Green fluorescing CaMPARI undergoes photoconversion (PC) to a red fluorescing protein only in the presence of intense 405 nm light and high Ca^2+^ levels. This conversion is irreversible, thus creating a temporal snapshot in a form of a ratiometric red/green output. CaMPARI is expressed exclusively in neuronal tissue due to a pan-neural promoter (*elavl3*, a.k.a. HuC) and is a direct read-out of relative neural activity. Initially, this innovation was not suitable for a screening tool as the photoconversion was not efficient enough. Thus, we have optimized the use of CaMPARI, such that the photoconversion is more efficient and practical for high throughput well-plate-formats. This allows our novel *in vivo* assay to measure neural activity and potentially screen large libraries of compounds. Our results were supplemented with a behavioral assay obtained from the same larvae, which offers an independent proxy measure that is related to neural activity.

Here, we investigated the effects of the two most abundant cannabinoids found in the plant, THC and CBD, on neural activity and characterized CB1R and CB2R *in vivo* on the CBD effects in zebrafish. The effects of early THC and CBD exposure from immediately after egg fertilization until the end of gastrulation, at 10 hour post-fertilization (hpf), were monitored later in embryonic development, at 4 and 5 days post-fertilization (dpf). The concentrations used in this study mimic plasma levels of human subjects with high cannabis consumption^35–37^, but we do have to consider that the molecules have to pass from the waterbath through the chorion into the plasma of the zebrafish embryo. THC and CBD were studied individually and then together to understand the impact of individual compounds by themselves and to partially mimic the cannabis available for recreational purposes. Further, we characterized the cannabinoid receptors involved using established receptor inhibitors for CB1R and CB2R. This was important, because with many receptors being linked to cannabinoids *in vitro*, additional functional *in vivo* evidence would be of great value.

## Results

### High-throughput assessment of CaMPARI is a reliable metric of neural activity

In this study the objective was to gain insight in the effects of THC and CBD on neural activity during early embryonic development. We measured neural activity with CaMPARI. CaMPARI photoconverts irreversibly from a green to a red fluorescing protein only if user-applied 405 nm light application coincides with high calcium levels (Fig. 1a)^34^. Red and green fluorescing CaMPARI are quantified as ratio (referred here as “CaMPARI activity”). We expressed CaMPARI through a pan-neural promoter, *elavl3* exclusively in the CNS and thus as a read-out of neural activity (Fig. 1b). Here, we optimized the use of CaMPARI zebrafish for an automated high-content INCell2000 plate-reader (Fig.1c). CaMPARI transgene expression was most suitable for this platform at 4 dpf, which is a suitable balance of fluorescing intensity and a sufficiently developed brain.

**Figure 1:**
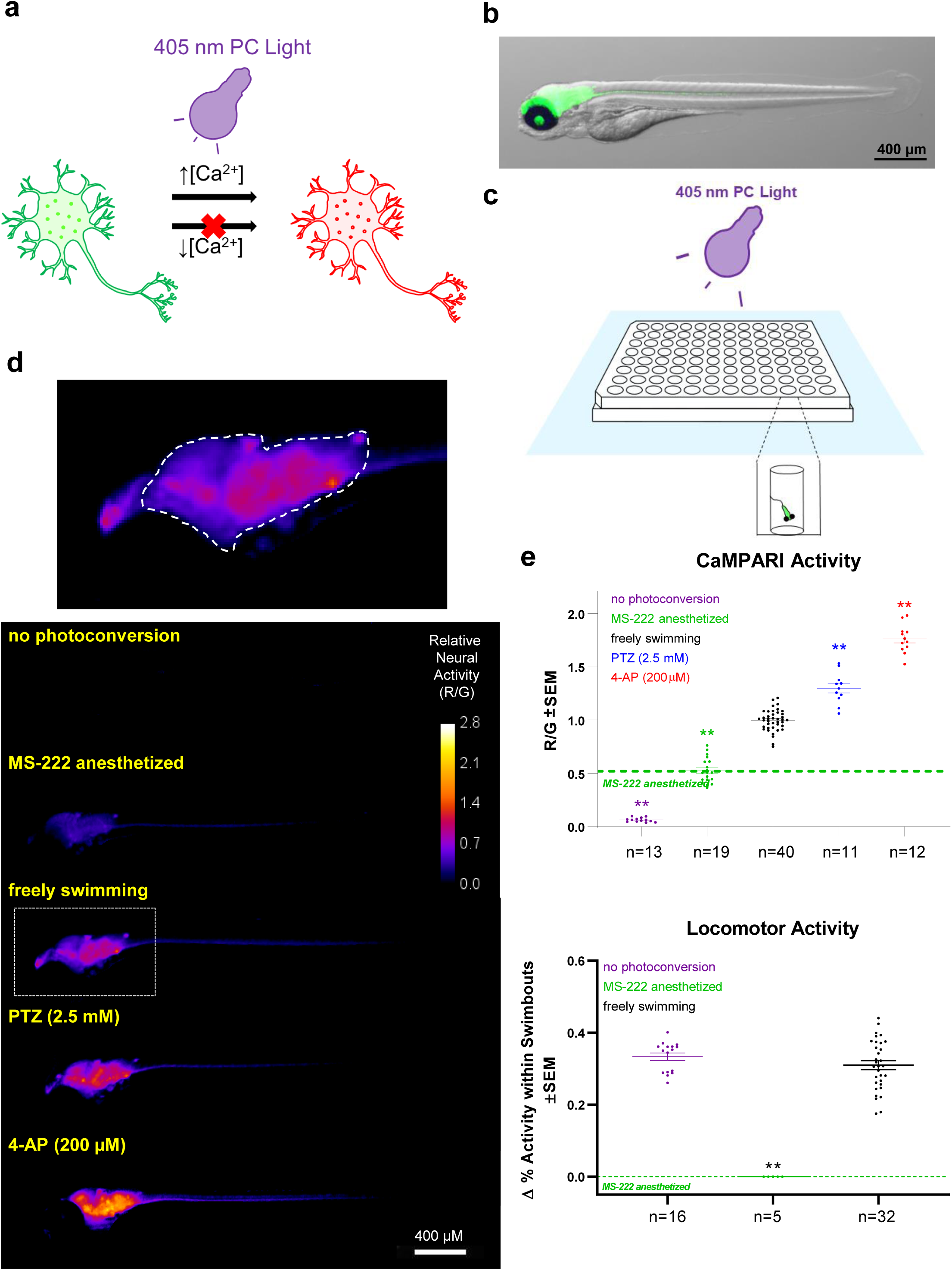
High-throughput quantification of neural activity in freely-swimming zebrafish larvae. (**a**) CaMPARI photoconverts from green to red fluorescing versions in the neuron only in the presence of both high intracellular calcium concentrations and a bright 405 nm light source. (**b**) Lateral view of green fluorescing CaMPARI merged with brightfield image shows exclusive expression in the CNS due to a pan-neural promoter (*elavl3*). (**c**) Larvae were transferred to 48 wells in the centre of a 96-well plate to ensure that the 405 nm LED Flood Array covers all larvae entirely. To minimize overheating, the plate was floating in a waterbath at 10 cm distant to the LED while CaMPARI was photoconverted (PC) by the LED Flood Array. (**d**) Lateral view of zebrafish with exemplar “CaMPARI Activity” heat maps. Heat maps show ratio of red/green (R/G) fluorescent output as indicated by the calibration bar and can be interpreted as relative neural activity. CNS regions with higher levels of neural activity translate to “hotter” pixels. Images were acquired using an automated INCell 2000 high-content microscope and reveal a reduction in neural activity when the larvae were anesthetized with MS-222 or an increase with convulsants, PTZ and 4-AP. Top, Enlarged lateral view from freely swimming larvae illustrating the optic tectum and hindbrain areas (dashed line) from which the R/G ratio was obtained. (**e**) Neural activity, inferred from the mean R/G ratio in optic tectum and hindbrain. CaMPARI activity is reduced to baseline in MS222-anaesthetized fish and photoconversion is undetectable when CaMPARI photoconverting light is omitted. Dashed green line represents mean for MS-222-anesthetized samples, which show a clear reduction in signal compared to freely swimming larvae and provides a baseline “near-zero” activity level for reference in subsequent experiments. Drivers of neural activity, PTZ and 4-AP induce a significant increase in CaMPARI activity. (**f**) Locomotor activity was measured using behavioural tracking software. PC light from the day prior at 4 days post-fertilization (dpf) did not affect locomotion at 5 dpf. MS-222 treatment abolished any swimbouts. Biological replicates are individual larvae = n. ** Significantly different from freely swimming samples, p<0.01.

CaMPARI activity was quantified and displayed as a heatmap consisting of the red/green ratio in fluorescing intensities. At 4 dpf, optic tectum and hindbrain regions of larvae showed a clear measureable output, whereas without photoconversion CaMPARI activity was nearly zero. To establish a baseline of near-zero neural activity, larvae were anesthetized with MS-222, and a significant reduction (p<0.01) in red/green ratio was obtained compared to freely swimming larvae. In contrast, drug-induced neural activity, with the established convulsants pentylenetetrazole (PTZ) or 4-aminopyridine (4-AP), showed a consistent increase in CaMPARI activity, even at concentrations considered to be minimal when inducing seizure (Fig. 1d,e)^38^.

We benchmarked our CaMPARI outputs against locomotor activity of zebrafish larvae, which is an established metric of neural activity^39^. We chose 5 dpf as an optimal developmental timepoint for analyzing locomotor activity, because younger larvae are largely inactive. Larvae were left in the same 96-well plate for assessing CaMPARI and locomotion. Exposing larvae to photoconversion with LED light the day before had no impact on the mean locomotor activity (Fig. 1f). Larvae anesthetized with MS-222 did not display any detectable swimbouts (Fig. 1f). The level of neural activity correlated with locomotor activity and will be presented later. Overall, these results show that CaMPARI, deployed in zebrafish larvae is a reliable high-throughput tool for measuring neural activity.

### CBD and THC reduce subsequent neural activity

We sought to assess how embryonic exposure to cannabinoids impacts upon subsequent neural activity later in development. Here, the doses of CBD and THC aligned with our previous work^13^, reflecting high cannabis consumption in humans. Comparisons of our dosage to humans requires various considerations: (i) blood plasma concentrations of THC can peak up to 0.25 mg/l while smoking a single cigarette^35^; (ii) the content of THC has increased in the past 20 years; and (iii) doses of intraperitoneally administered medical CBD can vary greatly, from 5-100 mg/kg, and daily maximum of 1500 mg/kg^36,37^. The current study uses up to 6 μg/mL of THC and 3 μg/mL of CBD. Absorption studies using Liquid Chromatography-Tandem Mass Spectrometry suggest that typically an estimated 0.1-10 % of toxic compounds will pass through the chorion to reach the embryo^40,41^.

We wanted to gain insight into the effects of CBD and THC separately (Fig. 2a), and therefore each was applied in a dose-response format. This validation was necessary as CaMPARI has not been used in studying cannabinoids. Compounds were added to the bath early in animal development at (0.5 hpf) and then washed out towards the end of gastrulation (10 hpf). CaMPARI was imaged at 4 dpf and locomotor activity was assessed at 5 dpf (Fig. 2b). Animals that were exposed to CBD at concentrations of 2 μg/ml and 3 μg/ml exhibited a dose-dependent reduction in CaMPARI activity (Fig. 2c,d). These reductions in CaMPARI output are substantial when compared against the ~50 % reduction we observed in anesthetized larvae, where little neural activity is expected. Coordinated with this, locomotion was also reduced starting at 1.5 μg/ml and was significant (p<0.01) at 3 μg/ml (Fig. 2e). THC had a similar effect as CBD and also reduced neural activity at higher doses, 4 μg/ml and 6 μg/ml, (Fig. 2f,g). Locomotor activity exhibited a more extensive reduction at 6 μg/ml (Fig. 2h). We compared neural activity and locomotion in the same individuals exposed to effective doses of cannabinoids and found significant correlations (Fig. S2; CBD r=0.52 (p<0.01) and THC r=0.71 (p<0.01)). Indeed, most larvae, that displayed reduced neural activity also showed reduced locomotion. Together, our findings show that both CBD and THC reduced neural activity when exposed early in development.

**Figure 2:**
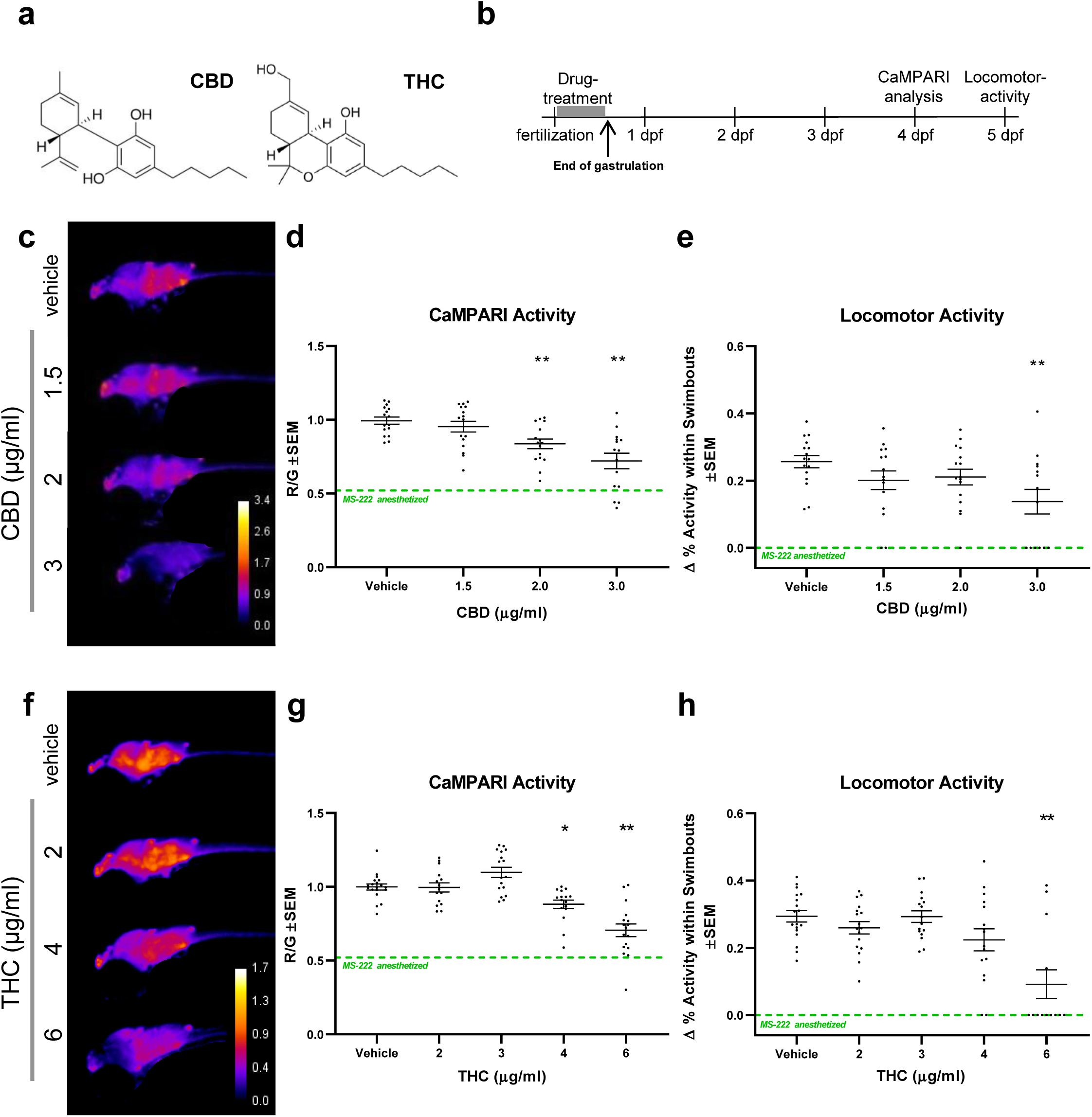
Early application of CBD and THC reduce subsequent neural activity and locomotor activity. (**a**) The chemical structures of key cannabinoids used in this study: CBD and THC. (**b**), Timeline of experimental set up where zebrafish embryos were treated with drugs between 0.5 and 1 hour post-fertilization (hpf) and washed out at 10 hpf towards the end of gastrulation. CaMPARI imaging was obtained at 4 days post-fertilization (dpf) and locomotor activity was tracked at 5 dpf. (**c**) and (**f**), Representative CaMPARI activity maps obtained from treated larvae and corresponding quantification and statistics shown in (**d**) and (**g**), respectively. (**e**) and (**h**), Locomotor activity of the same larvae. (**c**) CBD reduces neural activity and locomotor activity as shown in (**d**) and (**e**). (**f**) THC requires a higher does than CBD to reduce neural activity and locomotor activity as shown in (**g**) and (**h**). CaMPARI activity heat maps show ratio of R/G channels as indicated by the calibration bar, with higher ratios (hotter colours) representing greater neural activity. Green-dashed lines depict mean baseline (zero) value for MS-222 anesthetized samples (From Fig 1). Biological replicates are n=8-19. * is p<0.05; ** p<0.01 compared to the vehicle control (0.3 % MeOH in experiments with CBD and 0.6 % MeOH for THC).

### Antagonistic effects of CBD on neural activity is enhanced when combined with sub-effective doses of THC

Cannabis consumption during pregnancy exposes the fetus to THC and CBD in concert. In order to investigate whether CBD and THC have a combined effect that is different from either compound on its own, neural activity was measured when larvae were exposed to both cannabinoids, using the same timeline as the previous experiment (Fig. 2b). First, various sub-effective doses of CBD were applied from 0.5 to 1.5 μg/ml in the presence of 2 μg/ml THC (identified as sub-effective in Fig. 2). Preliminary dosage attempts revealed small, but significant differences with p<0.01 and p<0.05, respectively, on neural activity when combining 0.5 or 1.5 μg/ml CBD with 2.0 μg/ml THC (Fig. S3a,b). Locomotor activity also trended towards a dose-dependent reduction during concerted application of CBD and THC and was significant at 1.5 μg/ml with p<0.01 (Fig. S3c). Next, CBD and THC were added in a 1:1 ratio mixture at 1.0 and 2.0 μg/ml each. An enhanced antagonistic effect was revealed on neural activity when both CBD and THC were added at 2.0 μg/ml. CBD by itself at 2.0 μg/ml significantly (p<0.05) reduced neural activity (Fig. 3d,e) consistent with results in Fig. 2d, but CBD and THC together, further (p<0.01) reduced neural activity compared to CBD or THC alone (Fig. 3d,e). The combined effects of CBD and THC seemed synergistic in that CBD by itself at 2.0 μg/ml, although significant, had a relative small effect and THC by itself at 2.0 μg/ml had no effect. However, when CBD and THC at the same concentration were combined neural activity was reduced close to the mean of MS-222 anesthetized samples suggesting no to very little real neural activity. Locomotor activity was mainly affected by CBD with no additional reduction when compared to CBD and THC combined (Fig. 3f). No significant effect on neural activity or locomotor activity was obtained when 1.0 μg/ml of CBD and THC was applied (Fig. 3a-c). Due to the effect on locomotion, we also wanted to assess the integrity of neurons in connection to motor axons. Reticulospinal neurons in the hindbrain were stained with RMO44 antibody targeting NEFM, an established marker for neuronal damage^42,43^. Indeed, RMO44 immunostaining was further reduced when CBD and THC were combined at 2.0 μg/ml suggesting that both CBD and THC affect neuronal health (Fig. 4S). Together, these results suggest that CBD and THC are more potent in reducing neural activity when applied in combination.

**Figure 3:**
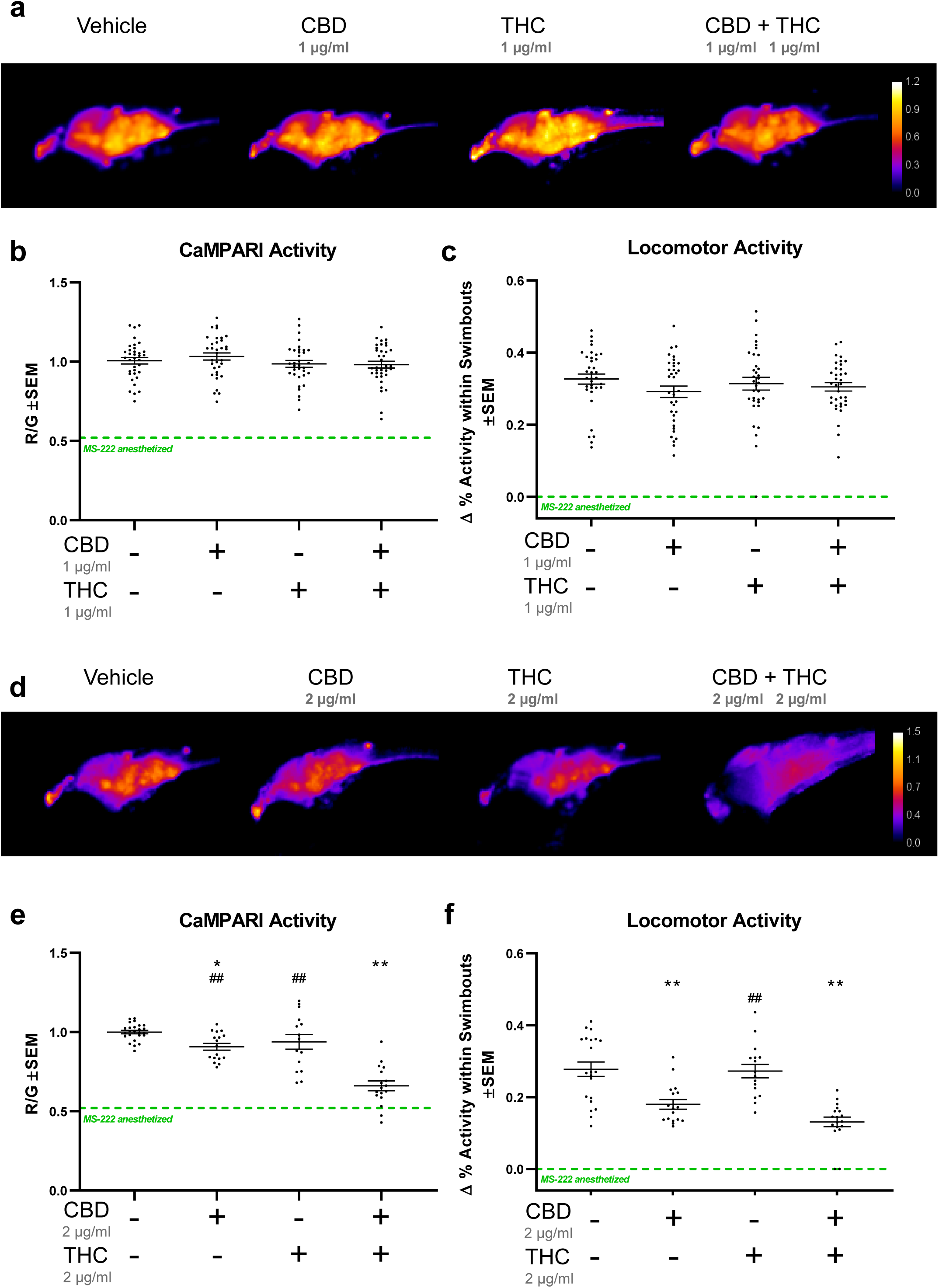
Antagonistic effect of CBD on neural activity is enhanced when combined with sub-effective doses of THC. Zebrafish larvae were exposed to a series of CBD and THC concentrations by themselves or in combination, which are mostly sub-effective, or in the case of CBD, were minimally effective. (**a**) and (**d**), Exemplar CaMPARI activity heat maps show an additive effect (minimizing neural activity) when exposed to 2 μg/ml (**d**) of each CBD and THC compared to CBD or THC by themselves (as plotted in (**e**)). When 1 μg/ml of 1:1 CBD and THC was applied, ratios show no additive effect as illustrated by quantifications and statistics in (**b**). (**c**) and (**f**), Locomotor activity from the same well at 5 dpf shows a clear reduction when CBD and THC is combined at 2 μg/ml each, but reduction mediated by CBD alone is almost as low as when combined with THC suggesting that CBD is the main component affecting locomotor activity. R/G is indicated by the calibration bar. Green-dashed lines depict mean values for MS-222-anesthetized samples (from Figure 1). Biological replicates are n=15-25. * compared to vehicle control (equal amount of MeOH in all experiments); ^#^ compared to CBD plus THC. One symbol is p<0.05; Two symbols is p<0.01.

**Figure 4:**
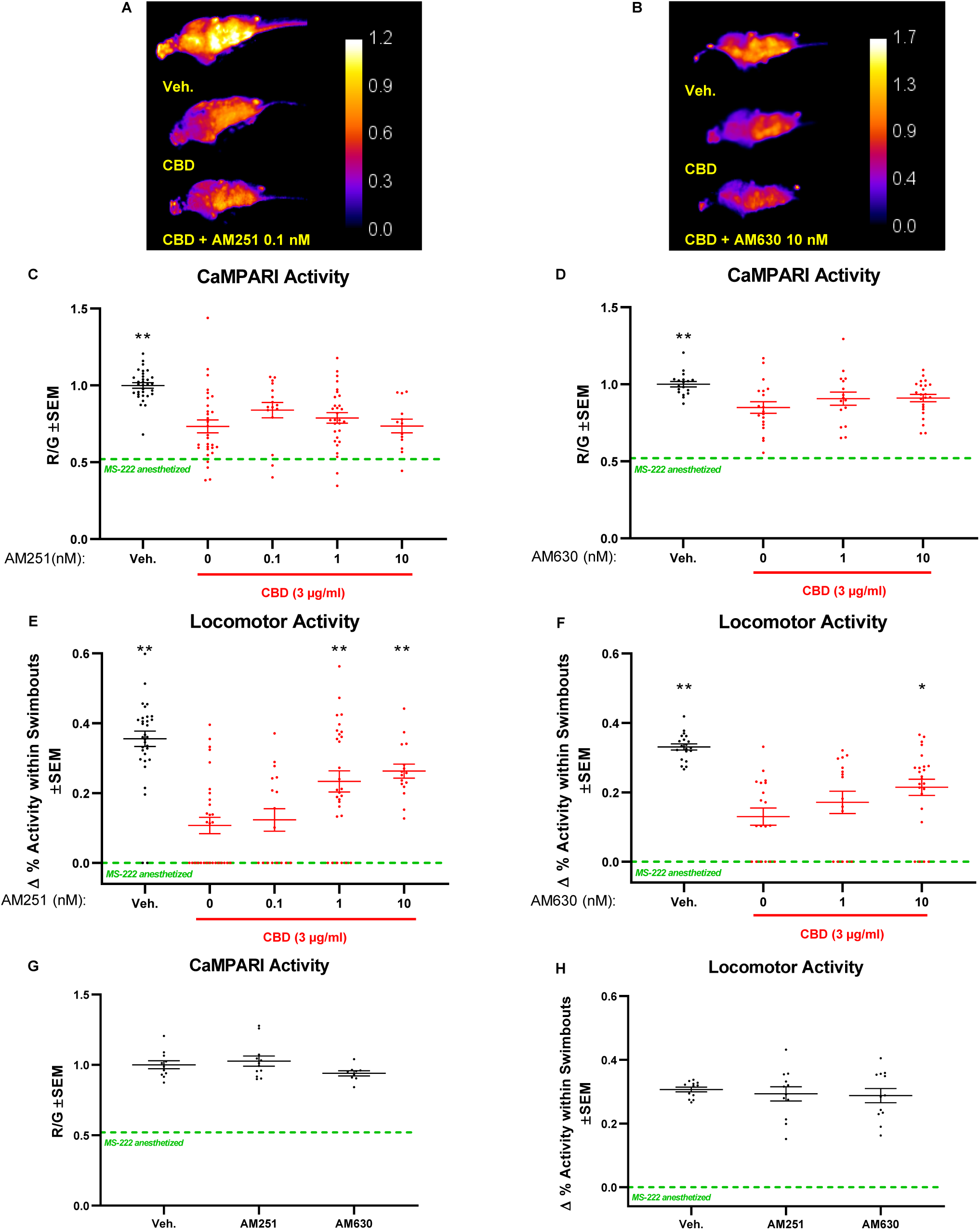
Cannabinoid receptor 1 (CB1R) and CB2R are both involved in CBD-mediated reduction of locomotor activity. Zebrafish larvae were exposed to CBD (3 μg/ml) with CB1R- and CB2R inhibitors, AM251 and AM630, respectively. (**a**) and (**b**), Exemplar CaMPARI heat maps showing R/G ratios obtained at 4 dpf. (**c**) and (**d**), Corresponding quantifications showing CBD mediated-reduction of neural activity is not fully rescued when CB1R is inhibited with AM251 or CB2R with AM630 in the nanomolar (nM) range. (**e**) and (**f**), Locomotor activity at 5 dpf shows a clear rescue when inhibiting either CB1R or CB2R with 0.1 nM AM251, or 10 nM AM630, respectively. (**g**) and (**h**), shows that applying AM251 or AM630 without CBD did not affect CaMPARI activity or locomotion. R/G is indicated by the calibration bar. All samples contained the same amount of vehicle DMSO (0.1 %) and MeOH (0.3 %) including the vehicle control (Veh.). Green-dashed lines depict mean values for MS-222 anesthetized samples. Biological replicates are n=13-30. * is p<0.05; ** p<0.01 compared to CBD.

### CB1R and CB2R are both required for CBD-induced reduction of locomotor activity

Some reports suggest that CBD mediates its actions through CB1- and CB2-receptors^25^. To test whether these two receptors are involved in the neuronal toxicity mediated by CBD *in vivo*, we deployed corresponding receptor inverse agonist AM251 and antagonist AM630. This pharmacological approach allowed the temporal specificity required for our experimental design. Both inhibitors appear to act with good specificity in zebrafish. Previous works support the specificity of CB1R and CB2R inhibitors used in this study^44,45^. In connection to CB2R, a recent study has shown that CB2 receptors are required for the action of AM630 on zebrafish behaviour. The impact AM630 had on wildtype larvae was not detectable when applied to *cnr2*^*-/-*^ mutant larvae, supporting drug specificity. Moreover, AM630 treated zebrafish showed very similar photo-dependent response compared to the CB2 knockout^44^. Similar in regards to CB1R, because knocking out CB1R and inhibiting CB1R with AM251 caused a similar response in rescueing locomotor activity when supressed with cannbinoid-receptor agonist, WIN55,212-2^45^. All three metrics measured (acclimatization, dark-adaption and recovery) for locomotion displayed arguably the same rescue-pattern in *cnr1*^*-/-*^ mutant and AM251 treated larvae^45^. Further, both inhibtors were used at nanomolar range in our study, which is considered modest and ensuring a level of specificity towards their respective receptors (AM251, IC50= 8 nM towards CB1R; AM630, IC50= 31.2 nM towards CB2R). As described previously, the experimental timeline is as depicted in figure 2b with CBD being added immediately after AM251 and AM630. When analyzing CaMPARI activity, our results revealed that the CBD-mediated reduction in neural activity could not be completely prevented with concentrations from 0.1 to 10 nM AM251 or AM630 in the presence of 3 μg/ml of CBD (Fig. 4a-d). In contrast, CBD-mediated reduction of locomotor activity was prevented with a concentration of AM251 as low as 1 nM (Fig. 4e) or with 10 nM of AM630 (Fig. 4f). In the absence of CBD, when using the maximum dose (10 nM) of AM251 or AM630, neither inhibitor had an effect (Fig. 4g,h). This finding excludes the possibility that the effects of AM251 or AM630 on locomotion were independent of CBD and rather specific towards CB1R and CB2R, respectively. Overall, this suggests a role for both CB1R and CB2R receptors in CBD-mediated effects on locomotor activity. In sum, inhibiting CB1R and CB2R can ameliorate the negative impacts of early CBD on late CNS function, at least with respect to locomotion.

## Discussion

In some jurisdictions, including Canada, cannabis has been legalized despite the lack of mechanistic understanding and health implications. Cannabis consumption has increased in the past decade^46,47^ and reports emerged associating prenatal cannabis consumption with stillbirths and autism spectrum disorder, a neurodevelopmental syndrome^48,49^. Yet, cannabis and CBD oil have been used by consumers to treat labor pain^50^. In animal models, such as rats and chicken, cannabis has been shown to have detrimental effects on developing embryos^10, 51^. Intriguingly, more recently, new evidence revealed that exposure to not only the more widely studied THC, but also CBD, during gastrulation impacts the overall development in zebrafish^13^. Interestingly, the effect of the two key cannabinoids, CBD and THC, were delayed, and observed much later in the development after the exposure had ended.

*In vitro*, CBD and THC can function through CB1R and CB2R, which are expressed in neuronal tissue^17–25^. Our goal was to specifically investigate the effects of CBD, how it might interact with the more widely studied THC during early development and characterize the impact on neural activity *in vivo* on a mechanistic level. For our study, we implemented CaMPARI transgenic zebrafish as a read-out of neural activity and supplemented this approach by measuring the locomotor activity of the same larvae. CaMPARI has proven to be a simple and economical neural integrator that can be optimized in a well-format using our animal model. We found that neural activity was reduced later in zebrafish development upon brief exposure of not only THC, but also CBD early in development. Surprisingly, CBD had similar impacts on neural activity at lower concentrations compared to THC. Notably, the impact of CBD was increased by sub-effective doses of THC. Our data quantifying locomotion, suggested that the CBD-mediated effect involves CB1R and CB2Rs.

We optimized CaMPARI such that this innovation can be used for large number of samples to screen compounds in future studies. Zebrafish larvae can be deployed in a 96-well set-up quantifying locomotion^39,52–56^. Nevertheless, locomotor activity is not a direct measurement of neural activity. Calcium sensors, such as the GCaMP series^57,58^, the more recently introduced GECO^59,60^, and CaMPARI, are more accurate. CaMPARI provides a temporal-snapshot of neural activity in the freely swimming animal, which can be anesthetized and imaged after the experiment^34^. Furthermore, the ratiometric red/green signals are imaged together and mitigate inter-individual differences in expression levels or focal plane. We optimized conditions for a well-plate set-up analyzing neural activity in the optic tectum and hindbrain. Unfortunately, less defined areas or specific neurons could not be quantified. Effects on motor axons will be detected by our behavioural assay. In general, there seems to be a good correlation between neural activity measured by CaMPARI and locomotor activity^61,62^. However, some findings may be more sensitive towards one assay vs the other, as described above. Furthermore, when interpreting our data as neural activity, it is important to consider that when anesthetized a clear red to green CaMPARI ratio can be still measured. This is unlikely due to autofluoresence because larvae that were not photoconverted yielded zero CaMPARI activity. We cannot exclude the possibility that due to longer photoconversion some CaMPARI has converted into the red fluorescence version even when anesthetized. It is well established that when anesthetized with MS-222, neurons are barely active^63,64^. Indeed, our behavioural assay shows no locomotion in the presence of MS-222 and we used this assay as a supplement in this study. We believe this method is a suitable technique to measure substantial changes in neural activity in the larger parts of the zebrafish CNS. For future studies in connection to epilepsy this innovation may be particularly helpful as some studies use convulstants, PTZ or 4-AP, to simulate seizure^62,65^. We found that both, PTZ and 4-AP, induce CaMPARI activity. Considering the advantages, this method outweighs its limitation and could serve for future studies, especially in connection with epilepsy.

We have considered several competing technologies prior to designing this study, in hopes that we can build upon and complement these potent approaches. Other than behavioural assays described above, the GCaMP series has been of interest^57,58^. However, the larvae have to be analyzed with high-resolution live microscopy during the experiment and movement of the larvae would add considerable variability. The assays would be data-intensive and technically challenging, where large sample sizes and different experimental groups are difficult to obtain (especially *in vivo*). Additionally, we were still interested in measuring outputs from freely behaving animals as constraining them for imaging likely impacts the results and is time-consuming. Other widely used markers for neural activity include the immediate-early gene *c-fos* and phosphorylation of ERK^65,66^. However, CaMPARI is more temporal specific due to immediate photoconversion. In addition, both markers are not only less tissue specific, but also respond to stress and other factors. The *elavl3* promoter that drives CaMPARI ensures CNS-specific expression. Overall, CaMPARI is a practical direct output of neural activity and thus the chosen innovation for this study.

Both CBD and THC have been implicated in embryonic development including abnormal changes to neurons. Delayed abnormalities after early exposure of key cannabinoids have only recently been investigated^13^. Here, early exposure of CBD and THC during the first 10 hours of embryonic development reduced neural activity when measured 4 days later, in a dose-dependent manner. These later effects may be due to abnormal neurodevelopment and/or the slow release of the cannabinoids from the tissue due to their lipophilic nature. Consistent with this finding, zebrafish larvae with a similar exposure (5-10 hpf) to cannabinoids have shown effects that could be due abnormal development (at 5 dpf). This was not exclusive to neurons, but included other changes such as morphological abnormalities, decreased survival rate, decreased mEPCs activity (may relate to muscle development), decreased heart rate and delayed hatching^13^. Considering the specific time-window of exposure (first 10 hours in development), an abrupt effect on the developmental program including neurons seems plausible. Interestingly, CBD seemed to impact neural activity at a lower concentration than THC, which also aligns with our previous study^13^. It is important to acknowledge that CBD consumption has been documented to help with seizures and cancer by reducing pain^67,68^. Nevertheless, our study suggests caution during pregnancies.

Considering that cannabis contains large proportions of both CBD and THC, we looked at the effects of these two compounds in combination and found a more potent effect in decreasing neural activity. Reticulospinal neurons in the hindbrain of the zebrafish control motor neurons during swimming^42,69^. In line with a reduction in locomotion, we observed reduced staining when exposed to both cannabinoids, suggesting that the reticulospinal neuronal integrity was compromised. However, this finding did not explain why locomotor activity was mainly influenced by CBD. This could be due in part that CBD has a greater effect at lower concentration than THC. The effect of CBD may be amplified by effecting motor neurons and potentially muscle development. Overall, these findings were surprising, because CBD was found to negate the effects of THC on CB1R^27^. One possibility is that at higher concentrations both, CBD and THC, affect the developmental program. This could explain the abnormal morphologies reported previously^13^. We have observed similar morphologies using higher concentrations of CBD and/or THC. Thus, it is of no surprise that abnormal morphologies impacted neural activity and locomotion. Nevertheless, because at 2 μg/mL we found a significant reduction in neural activity, but no morphological change was observed previously at that concentration^13^, it is unlikely that neural activity is affected by morphological changes alone. In short, we believe that although in physiological range, high concentrations of CBD and THC during embryonic development disrupt proper growth and neuron function with similar consequences and thus the effects of CBD and THC are same. This could result in overriding the negating effects of CBD. In contrast, at lower concentrations and in fully developed brains the impact of CBD and THC is different. Indeed, several receptors have been shown to have different affinities to THC and CBD^25^. A second possibility, therefore, is that at high concentration CBD and THC affect receptors other than CB1R and CB2R. This could explain why inhibiting CB1R and CB2R did not fully prevent CBD-mediated reduction of neural activity, as further discussed below.

At this stage, it is unclear what mechanism(s) account for the reduction in neural activity later in development when embryos are exposed briefly to CBD and THC in the first hours after fertilization. In this study, we were especially interested in the less widely studied CBD. Using HEK cells several receptor candidates have been identified showing to interact with CBD^25^. A recent study supports a functional role of CB1R and CB2R in zebrafish^32^. Here, by inhibiting CB1R, and CB2R, we found that the reduction in locomotor activity mediated by CBD was partially prevented at nanomolar concentrations of the inhibitors. This suggests that, mechanistically, CBD acts through both receptors. Previous studies have implied that CBD may bind to both CB1R and CB2R *in vitro*^25^ and our study is in support that these interactions may be important *in vivo*. In line with our findings, CBD has been shown to cause abnormal craniofacial and brain development in mice and zebrafish, which seems to be mediated through CB1R and the hedgehog signaling pathway^16^. Whether CB2R can mediate its effects through the same pathway remains to be determined. An explanation to why inhibiting CB1R and CB2R did prevent the maximum effect of CBD, which is delayed, could relate to the fact that the CB1R and CB2R are expressed very early in development and found on stem cell progenitor cells^61^. As mentioned above, an early effect may impact the developmental program such that changes persist until later, even if cannabinoids are no longer present in the circulation. Such changes may not be specific to the CNS as CB receptors are not only expressed in neuronal progenitor cells, but in stem cells forming other types of tissues^70^. This could explain our previous findings where CBD affected heart-rate and overall morphology after 5 dpf when exposed only briefly during gastrulation^11^. The CBD-mediated reduction in neural activity was not prevented when either CB1 or CB2Rs were inhibited. While the reason for this is unclear, it could relate that the effects of CBD are larger outside the optic tectum and the hindbrain, thus more visible when quantifying locomotion. As described above, our CaMPARI assay in well-format is not suitable to resolve motor axons in detail, which have been shown to be affected previously^13^. It is also possible that other receptors such as TRPV1 are involved^26^, but this needs further investigation.

Overall, both CBD and THC exposure during embryonic stages had a negative impact on later neural activity, which was additive when combined. Together, this could mean that the impact of cannabis on early development is higher than using isolated compounds and profound. Our findings also support an *in vivo* mechanism of CBD functioning through CB1R and CB2R. We believe this study opens up a new path for investigating detailed mechanisms, including other receptors, in specific neurons that are affected by cannabinoid toxicity early in development and persist. The accumulating evidence for prenatal cannabis consumption having negative consequences on neurodevelopmental disorders in the offspring^3^ has to be considered. As the recreational use is becoming legalized and socially acceptable, more studies are warranted to not only fully understand the impact on human development, but also the mechanisms of the diseases.

## METHODS

### Animal Ethics and exposure to drugs

Zebrafish maintenance was approved by the Animal Care & Use Committee: Biosciences at the University of Alberta and operated under the guidelines of the Canadian Council of Animal Care. The fish were maintained in the University of Alberta fish facility at 28°C under a 14/10 light/dark cycle as previously described^71^ and all authors complied with ARRIVE (Animal Research: Reporting of In Vivo Experiments) guidelines^72^.

CaMPARI transgenic zebrafish were outcrossed with *Casper* strains not carrying the transgene. For experimental set up, embryos were randomly collected and placed in egg water (60 μg/ml Instant Ocean) in groups of 20 embryos as early as practically possible within 0.5 hpf. Stock solutions of CBD and THC were obtained from Sigma at 1 mg/ml dissolved in methanol. Receptor inhibitors AM251 (Selleck Chemicals, Houston, TX, USA) and AM630 (Adooq Bioscience, Irvine, CA, USA) were dissolved in DMSO. Accordingly, zebrafish embryos were treated with vehicle or compounds as close to 1 hpf as practically possible and removed towards the end of gastrulation (10 hpf). Within experiments all treatments contained the same amount of methanol or DMSO, specific percentages are indicated in the figure legends. Compounds were removed with several washes and egg water was replaced every day until CaMPARI analysis. Not all larvae carried the transgene and were screened for green fluorescence. An increase in mortality and abnormal morphology was observed when treated with higher doses of CBD or THC, as reported previously^13^.

### Engineering CaMPARI transgenic zebrafish for integrative calcium imaging

Zebrafish expressing CaMPARI, i.e. the Tg[*elavl3*:CaMPARI (W391F+V398L)]^ua3144^ line were generated as described previously^61,62^. We re-derived CaMPARI fish in response to a federal moratorium limiting zebrafish import into Canada^73^.

### CaMPARI photoconversion

For photoconversion, bright green Tg[*elavl3*:CaMPARI (W391F+V398L)]^ua3144^ larvae at 4 dpf were placed in a 96-well plate containing 150 μl egg water (made as previously decribed^71^) and acclimatized for 2 h. The central 48 wells of the plate were exposed to 405 nm LED array (Loctite) for 300 sec at a distance of 10 cm. LED array illuminated the plates entirely and evenly. The plates were floating in water at room temperature to ensure that the larvae did not overheat. For imaging, the larvae were anesthetized in 0.24 mg/mL tricaine methanesulfonate (MS-222, Sigma) following photoconversion. MS-222 was washed out and zebrafish were rested for a day prior analysis for locomotor activity.

### CaMPARI analysis

The ratio of red to green fluorescent emissions (red photoconverted CaMPARI in ratio to green CAMPARI) was interpreted as relative neural activity, as previously defined^34^. Images were objectively obtained with an InCell 2000 microscope (GE Healthcare, US). 96-well plates were run using FITC (1 second exposure) and dsRed (2 second exposure) channels. Blinded experimentalists used ImageJ to quantify the fluorescence mean of the red (dsRed) and green (FITC) channels. The area of the optic and hindbrain area were quantified as those regions were most consistent in the view. A mask was applied to exclude noise outside of the CNS.

### Quantifying Locomotor activity

The same zebrafish larvae being placed in the 96-well plate the day prior for CaMPARI analysis were objectively quantified for locomotion using Basler GenICaM (Basler acA 1300-60) scanning camera (75-mm f2.8 C-mount lens) and EthoVision^®^ XT-11.5 software by Noldus (Wageningen, Netherlands) as described previously^74^. Briefly, larvae were acclimatized for 3 h and one hour of movement was recorded from each individual larvae in each well. Beneath the plate an infrared backlight source was located and the scanning camera above. Activity was defined here as % pixel change within well between frames (recordings were at 25 frames per second) and quantified with EthoVision^®^ XT-11.5 software. This set-up allowed to see whether there is a correlation between level of neural activity and the the mean activity % within a swimbout (swimbout size). Swimbout size was shown to increase in response to the convulsant PTZ previously^52^ and here it can be a useful marker for neural activity. Transparent *Casper* strain zebrafish^75^ were unable to be tracked as ususal with Noldus. Therefore, quantification of swimbouts provided an additional advantage. Clear movement was detected above the noise treshold due to movement of the pigmented eyes. Although the % swimbout is small, reliable measurements were obtained. Only clear swimbouts were quantified, which were set above a treshold of 0.1 % of activity. Figure 1f shows that measuring the mean activity of a swimbout is a useful tool in obtaining output that is clearly diminished in presence of MS-222. In addition, the output also shows sensitivity in freely swimming larvae.

### RMO44 staining

The zebrafish used in this experiment were of the Tuebingen Longfin strain. Set-up was similar as described above. Following the experiment, larvae were stained with anti-RMO44 (Thermo Fisher Scientific, Waltham, MA, USA) and imaged using a Zeiss LSM 710 confocal microscope (Oberkochen, Germany) as described elsewhere^76^.

### Statistics

GraphPad Prism Software (Version 7, GraphPad, San Diego, CA) was used to analyze statistics from data obtained from our CaMPARI assay or tracking locomotor activity. All the data were presented as mean ±SEM (standard error of the mean). Statistical significance of p<0.05 was determined using a non-parametric t-test between two groups followed by a Mann-Whitney analysis, if approriate. Multiple groups were compared using One-Way ANOVA with Dunnett’s multiple comparisons test. Pearson r from correlation analysis was also determined where indicated.

## ACKNOWLEDGEMENTS

This study was supported by grants to W.T. Allison from Alberta Prion Research Institute-Alberta Innovates BioSolutions and the Alzheimer Society of Alberta and the Northwest Territories, and to D.W. Ali from the Natural Sciences and Engineering Research Council of Canada and Alberta Innovates mCannabis. L.F. Locskai was supported by undergraduate studentships from Natural Sciences and Engineering Research Council of Canada and from Alberta Innovates Health Solutions. R. Kanyo was supported by a SynAD postdoctoral fellowship funded via Alzheimer Society of Alberta and Northwest Territories through their Hope for Tomorrow program and the University Hospital Foundation. We would like to also extend our thanks to members of the Allison and Ali lab. In addition, we would also like to acknowledge the Sciences Animal Support Services at the Unversity of Alberta.

## AUTHOR’S CONTRIBUTIONS

RK, MRA, WTA, DWA contributed to the experimental design. RK, MRA, LFL, DDB, AMO, performed the experiments and analyzed the data. RK, MRA and DWA interpreted the results. RK, WTA and DWA wrote the manuscript. All authors read and approved the manuscript.

---

**Figure S1: Supplementary figure accompanying Fig.2.** Re-plotting data to illustrate that CBD shows an effect at lower concentration than THC. Quantifications of CaMPARI activity (left) or locomotor activity (right) showing a statistical difference between CBD and THC at 4 or 5 days post-fertilization when adding 3.0 μg/ml of either cannabinoids. Green-dashed lines depict mean values for MS-222 anesthetized samples. Biological replicates are n=14-28. Statistics using unpaired t-test shows ** p<0.01 compared to CBD.

**Figure S2: Supplementary figure accompanying Fig.2.** Re-plotting data from groups that were treated with vehicle, CBD (3 μg/ml) or THC (6 μg/ml) to assess correlation between CaMPARI activity vs. Locomotor activity within the same larvae. There is a significant positive linear correlation between neural activity and locomotor activity when corresponding vehicle controls (black circles) were plotted with either CBD (**a**; red circles) or THC (b; blue circles). Green-dashed lines depict mean values for MS-222 anesthetized samples.

**Figure S3: Supplementary figure accompanying Fig.3.** Additive effects in reducing neural activity are also obtained at lower CBD concentrations while THC concentration is kept constant at 2 μg/ml. (**a**) Exemplar CaMPARI activity heat maps obtained at 4 days post-fertilization show an effect at 0.5 and 1.5 μg/ml of CBD when combined with 2 μg/ml THC (blue) compared to CBD alone (red), as illustrated by quantifications in (**b**). (**c**) Locomotor activity from the same well at 5 dpf. R/G as indicated by the calibration bar. Green-dashed lines depict mean values for MS-222 anesthetized samples. Biological replicates are n=14-28. * is compared to CBD; ^#^ is compared to Vehicle. One symbol is p<0.05; two symbols are p<0.01.

**Figure S4: Supplementary figure accompanying Fig.3.** RMO44 staining of reticulospinal neurons in the hindbrain shows a decrease in fluorescence intensities when combining CBD and THC at 2 μg/ml at 5 dpf compared to when CBD or THC was added alone. Images show biological replicates (n=3), left to right, from corresponding treatment group, top to bottom.

