## Supplemental Figures for "Medium-throughput zebrafish optogenetic platform identifies deficits in subsequent neural activity following brief early exposure to cannabidiol and Δ-9-tetrahydrocannabinol"

#### Fig. S1

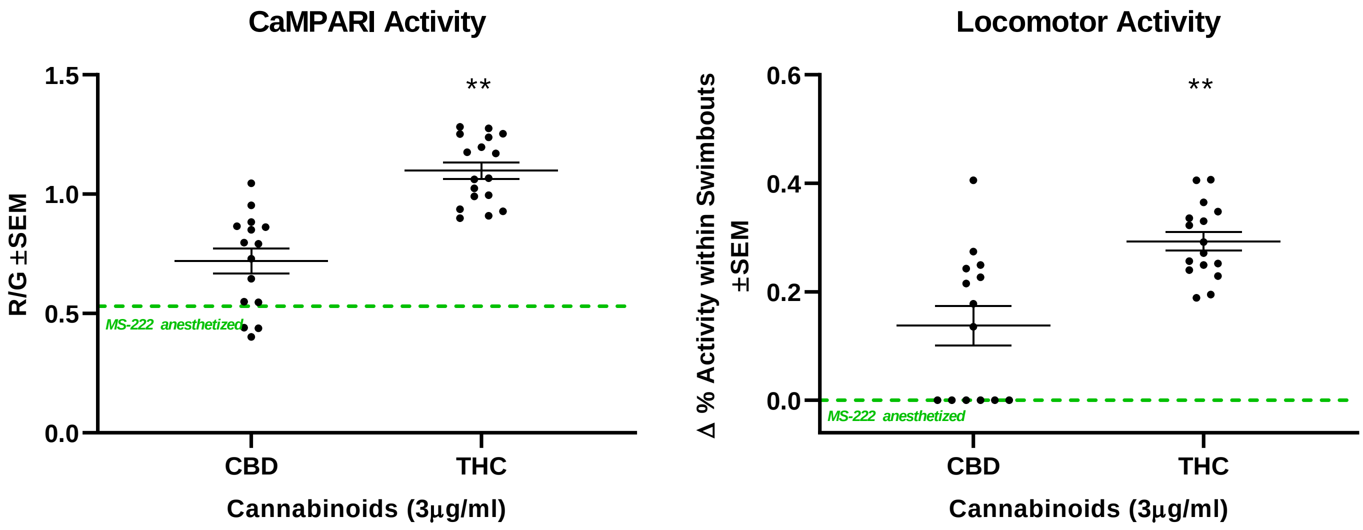
CaMPARI Activity
R/G ±SEM
1.5
1.0
0.5
0.0
**
MS-222 anesthetized
CBD
THC
Cannabinoids (3µg/ml)
Locomotor Activity
Δ % Activity within Swimbouts ±SEM
0.6
0.4
0.2
0.0
**
MS-222 anesthetized
CBD
THC
Cannabinoids (3µg/ml)

Fig. S2

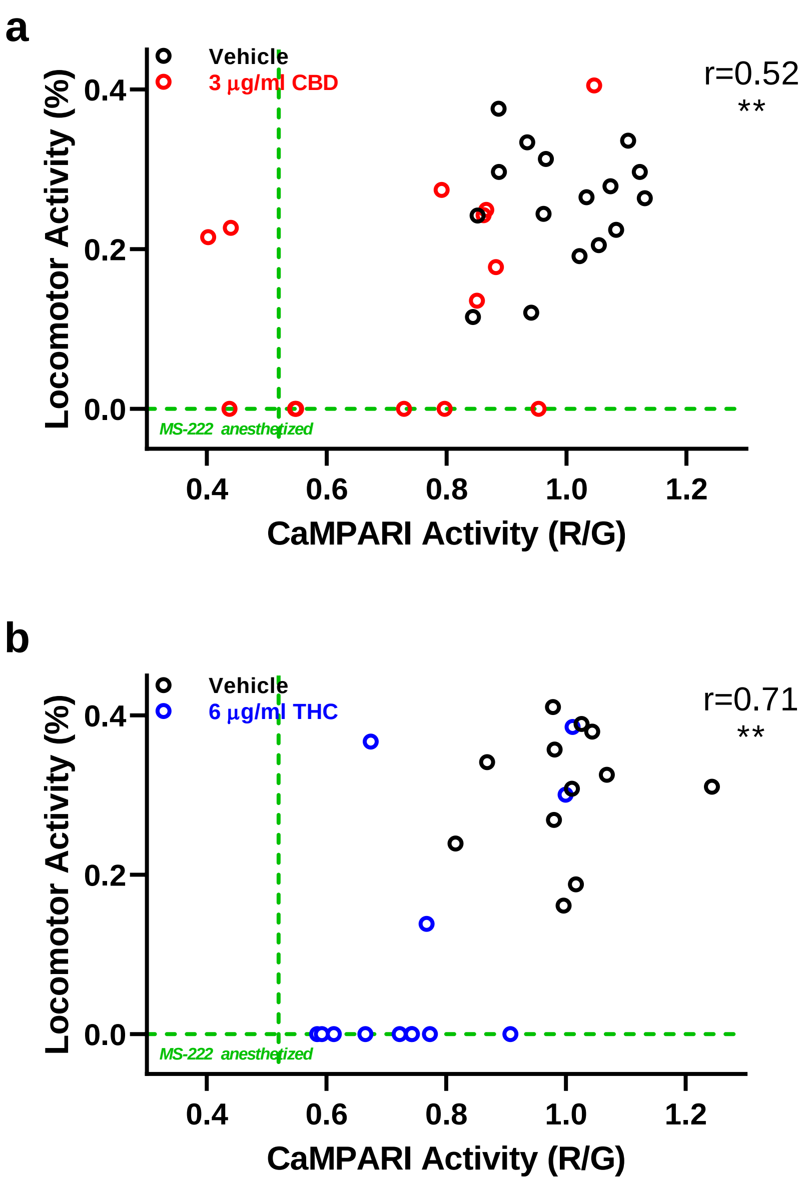
a
Vehicle
3 µg/ml CBD
r=0.52
**
Locomotor Activity (%)
0.0
0.2
0.4
MS-222 anesthetized
0.4
0.6
0.8
1.0
1.2
CaMPARI Activity (R/G)
b
Vehicle
6 µg/ml THC
r=0.71
**
Locomotor Activity (%)
0.0
0.2
0.4
MS-222 anesthetized
0.4
0.6
0.8
1.0
1.2
CaMPARI Activity (R/G)

**Fig. S3**

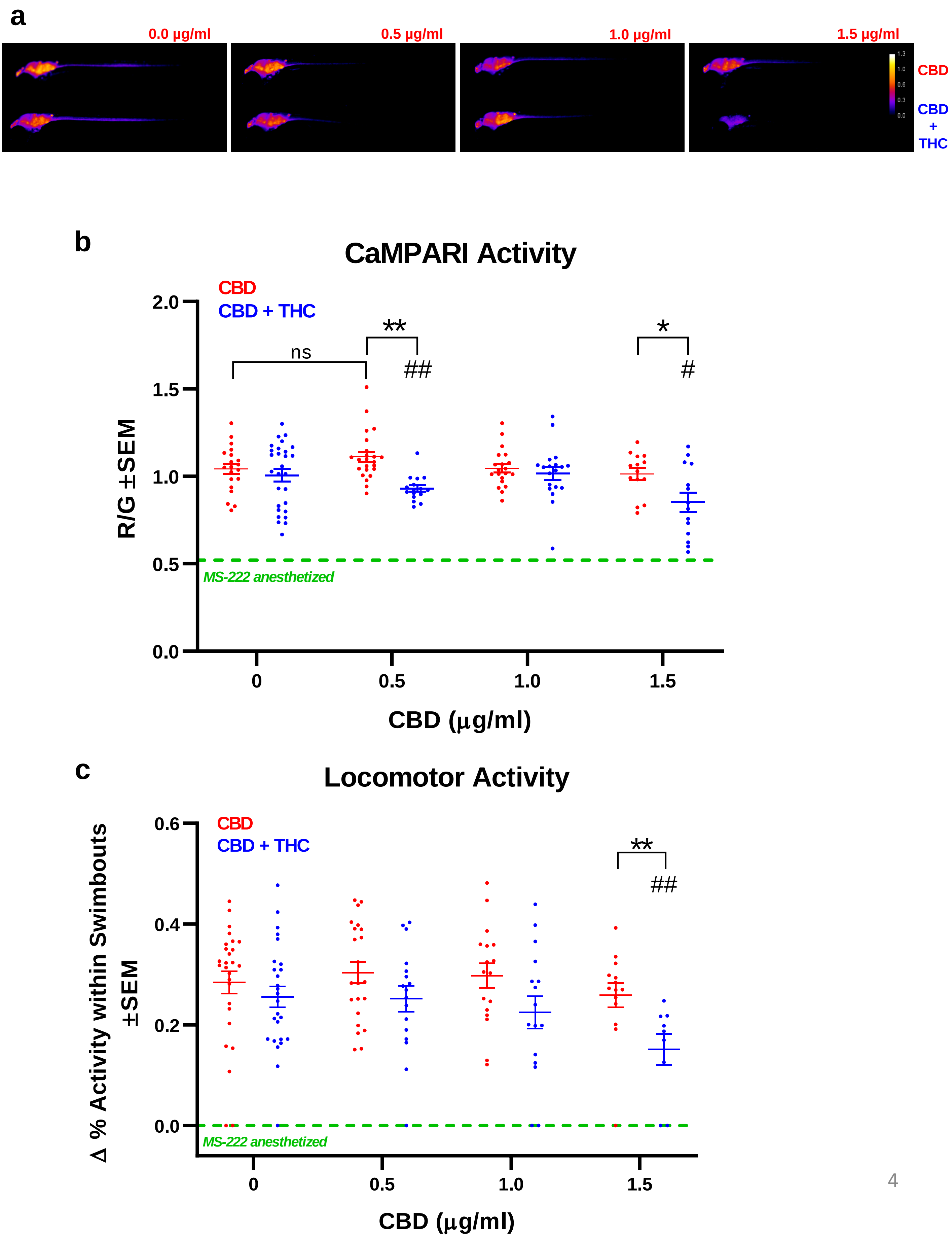
a
0.0 µg/ml
0.5 µg/ml
1.0 µg/ml
1.5 µg/ml
1.3
1.0
0.6
0.3
0.0
CBD
CBD + THC
b
CaMPARI Activity
CBD
CBD + THC
2.0
1.5
1.0
0.5
0.0
ns
**
##
*
#
MS-222 anesthetized
R/G ±SEM
0
0.5
1.0
1.5
CBD (µg/ml)
c
Locomotor Activity
CBD
CBD + THC
0.6
0.4
0.2
0.0
**
##
MS-222 anesthetized
Δ % Activity within Swimbouts ±SEM
0
0.5
1.0
1.5
CBD (µg/ml)

### Fig. S4

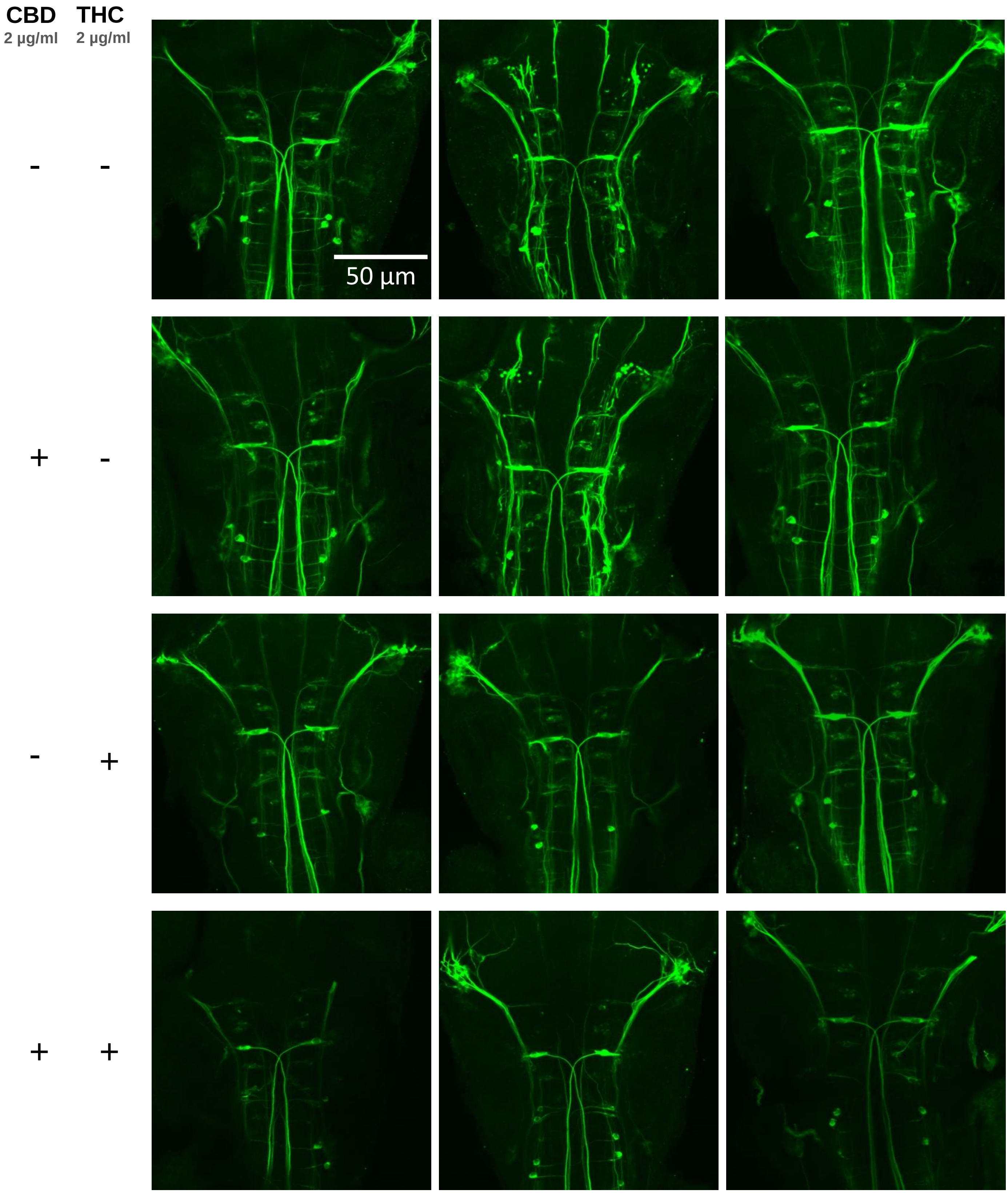
